# On the relationship between instantaneous phase synchrony and correlation-based sliding windows for time-resolved fMRI connectivity analysis

**DOI:** 10.1101/179820

**Authors:** Mangor Pedersen, Amir Omidvarnia, Andrew Zalesky, Graeme D. Jackson

## Abstract

Correlation-based sliding window analysis (CSWA) is the most commonly used method to estimate time-resolved functional MRI (fMRI) connectivity. However, instantaneous phase synchrony analysis (IPSA) is gaining popularity mainly because it offers single time-point resolution of time-resolved fMRI connectivity. We aim to provide a systematic comparison between these two approaches, on both temporal and topological levels.

For this purpose, we used resting-state fMRI data from two separate cohorts with different temporal resolutions (45 healthy subjects from Human Connectome Project fMRI data with repetition time of 0.72 s and 25 healthy subjects from a separate validation fMRI dataset with a repetition time of 3 s). For time-resolved functional connectivity analysis, we calculated tapered CSWA over a wide range of different window lengths that were temporally and topologically compared to IPSA.

We found a strong association in connectivity dynamics between IPSA and CSWA when considering the absolute values of CSWA. This association peaked at a CSWA window length of ∼20 seconds, irrespective of the sampling rate of the underlying fMRI data. Narrow-band filtering of fMRI data (0.03-0.07 Hz) yielded a stronger relationship between IPSA and CSWA than wider-band (0.01-0.1 Hz). On a topological level, time-averaged IPSA and CSWA nodes were non-linearly correlated, mainly because nodes with strong negative correlations (CSWA) displayed high phase synchrony (IPSA).

Our results suggest that IPSA and CSWA provide comparable characterizations of time-resolved fMRI connectivity for appropriately chosen window lengths. Although IPSA requires narrow-band fMRI filtering, we recommend the use of IPSA given that it does not mandate a (semi-)arbitrary choice of window length and window overlap. A MATLAB code for calculating IPSA is provided.

## 1. Introduction

Functional MRI (fMRI) connectivity has become a well-established paradigm of interrogating spatial and topological aspects of human brain function, in health and disease (Biswal et al., 1995; Bullmore and Sporns, 2009; Fornito et al., 2015). A recent trend in the field of macroscopic neuroimaging has been to treat fMRI connectivity data as a time-resolved function rather than averaging the functional connectivity over the course of the scan (Hutchison et al., 2013; Thompson and Fransson, 2015).

Correlation-based sliding-window analysis (CSWA) is the most commonly used method to calculate time-resolved fMRI connectivity (Allen et al., 2012; Hudetz et al., 2015; Kaiser et al., 2016; Rashid et al., 2016; Shakil et al., 2016; Tagliazucchi and Laufs, 2014; Yu et al., 2015; Zalesky et al., 2014; see also Preti et al. 2016, for a comprehensive overview of fMRI based CSWA studies). This approach entails correlating the fMRI signals between distinct brain regions within a pre-defined number of fMRI time points that form a ‘window’. The window is successively shifted over time to generate a time-resolved measure of interregional functional brain connectivity, reflecting presumed temporal changes associated with spontaneous brain network activity. Instantaneous phase synchrony analysis (IPSA) has gained popularity recently (Córdova-Palomera et al., 2016; Demirtaş et al., 2016; Glerean et al., 2012; Omidvarnia et al., 2016; Pedersen et al., 2017b, 2017a; Ponce-Alvarez et al., 2015) and does not mandate the often-arbitrary choice of window length and window overlap inherent to CSWA. Furthermore, IPSA ensures that time-resolved functional connectivity is characterised at the same temporal resolution as the input (narrow-band) fMRI signal, whereas the resolution of CSWA is largely governed by the choice of window length.

Our main objective in this paper was to provide a comprehensive yet objective comparison between IPSA and CSWA from a temporal and topological perspective. Glerean et al. (2012) provided evidence suggesting that IPSA and CSWA are temporally inter-related. Here, we build on this important previous study by evaluating several key aspects of time-resolved fMRI connectivity. These included the effects that *i)* CSWA window length, *ii)* CSWA negative correlations, *iii)* fMRI sampling rate and *iv)* fMRI band-pass filtering have on the relationship between IPSA and CSWA.

While IPSA and CSWA are ubiquitous, it is important to recognize that alternative methods to map time-resolved functional brain connectivity have been developed, including temporal independent component analysis (Smith et al., 2012), model-based approaches (Lindquist et al., 2014), time-frequency coherence analysis (Chang and Glover, 2010) and change-point detection methods to identify stationary time segments (Cribben et al., 2013). These methods were not considered in this paper.

The remainder of the paper is structured as follows: Firstly, we introduce our two resting-state fMRI datasets and provide an overview of the mathematical properties of IPSA and CSWA. We then present our results regarding the association in connectivity dynamics between IPSA and CSWA, on a whole-brain level and also between specific connection pairs. The topological association between IPSA and CSWA was then considered. Lastly, we discuss the wider implications of using IPSA instead of CSWA as a measure of time-resolved fMRI connectivity, and we also provide MATLAB code for calculating IPSA.

## 2. Methods and Materials

### 2.1 fMRI cohort

We used resting-state fMRI data acquired in 45 healthy subjects (age range = 22-35 years) from the Human Connectome Project (HCP - Van Essen et al. 2013) (left-right encoded, first session scans of the 3T test-retest dataset - Young Adults 1200 subject data release available at: https://db.humanconnectome.org/) with 72 slices (each slice was 2 mm thick) with a repetition time (TR) of 0.72 seconds. Echo time was 58 milliseconds, flip angle was 90° and voxel size was 2×2×2 mm. In total, 14.4 minutes (1200 time points) of multiband fMRI data was obtained per subject.

### 2.2 fMRI validation cohort

To ensure that our findings were reproducible, we also included a separate validation cohort comprising 25 healthy subjects (mean age = 25.4 years, ± 5.2 standard deviation). Resting-state fMRI data were recorded using a 3T Siemens Skyra MRI system (Erlangen, Germany) with 44 slices of 3 mm thickness, TR of 3 seconds, echo time of 30 milliseconds, flip angle of 85° and voxel size of 3×3×3 mm. Ten minutes (200 time points) of fMRI data was obtained per subject. All subjects gave written informed consent to participate in the study and it was approved by the Austin Health Human Research Ethics Committee, Austin Hospital, Melbourne, Australia.

### 2.3 fMRI pre-processing

FMRI pre-processing was done separately for our two cohorts using the SPM12 toolbox (Friston et al., 2011) in the MATLAB 2016b environment (MathWorks Inc., Natick, Massachusetts, United States). The fMRI pre-processing parameters were identical for both datasets apart from slice timing correction, which was only performed on the validation cohort (Fig. 1). This is because it is non-trivial to slice-time correct multi-band fMRI data (Glasser et al., 2013). The fMRI data was realigned and co-registered to the participant’s own T1 weighted image that was skull-stripped using FSL BET (Smith, 2002). Group-level segmentation of grey matter, white matter and corticospinal fluid was done using DARTEL (Ashburner, 2007). We denoised the fMRI data using 24 motion realignment parameters (Friston et al., 1996) and CompCor (Behzadi et al., 2007). CompCor removes the first five principal components associated with white matter and cerebrospinal fluid fMRI signals by using spatial masks obtained in the previous segmentation step. Each subject’s fMRI data was then spatially normalised into Montreal Neurological Institute (MNI) space using group template estimates from DARTEL. We did not spatially smooth the pre-processed fMRI datasets to prevent spurious correlations between adjacent brain nodes and we did not remove fMRI time-points associated with head movements in order to avoid temporal discontinuity in the fMRI time-series.

**Figure 1:**
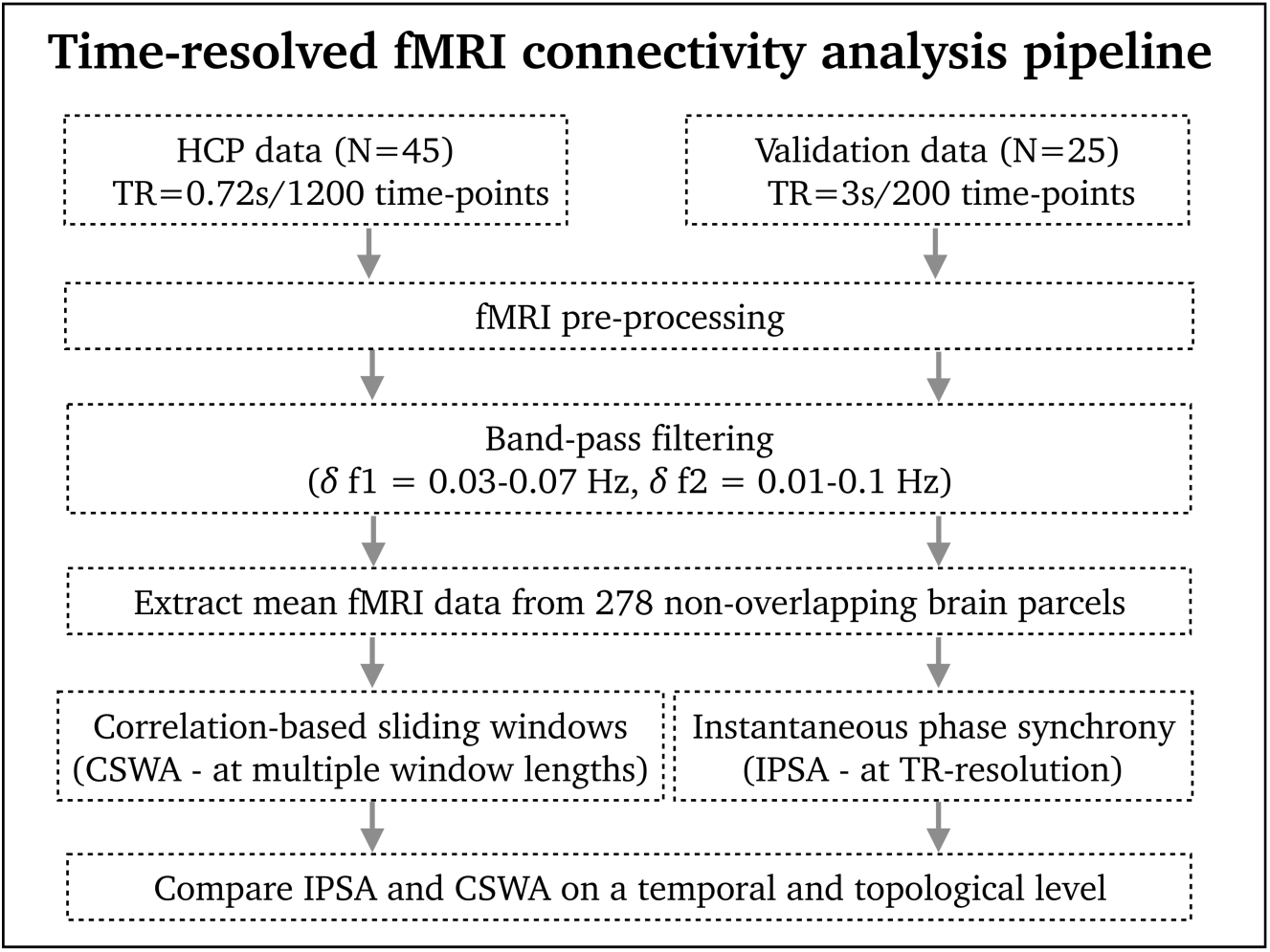
An overview of the adapted analysis paradigm in this study.

An important aspect of our time-resolved fMRI connectivity framework was band-pass filtering of the data. We filtered the fMRI time-series in two different ways: *i)* a narrow band-pass filter between 0.03 and 0.07 Hz which is thought to satisfy Bedrosian’s requirement for phase synchrony analysis (Bedrosian, 1963; Glerean et al., 2012; Omidvarnia et al., 2016; Ponce-Alvarez et al., 2015), and *ii)* a wider band-pass filter between 0.01 and 0.1Hz which is commonly used in correlation-based fMRI studies (Biswal et al., 1995; Fransson, 2005).

We calculated time-resolved fMRI connectivity (IPSA and CSWA) between 278 non-overlapping brain parcels (*N = 278*) using the average fMRI signal across all voxels in each node. This brain parcellation mask was previously defined on the basis of functionally homogenous brain regions across 79 healthy brains (see Shen et al., 2013, for more information). Our primary results were not driven by this particular parcellation mask as a lower resolution parcellation mask consisting of 90 nodes (Tzourio-Mazoyer et al., 2002) elicited near identical results to the original analysis with 278 nodes (see Supplementary materials 1).

### 2.4 Instantaneous phase synchrony analysis (IPSA)

IPSA was calculated similarly to previous studies (e.g., Pedersen et al., 2017b and Ponce-Alvarez et al., 2015) where the Hilbert transform (Mormann et al., 2000) was used to obtain phase information between all possible node-wise fMRI time-series. Let ***Y*** be a two-dimensional matrix of size *N × T* including average fMRI time-series across *N* brain parcels where *T* is the number of fMRI volumes. 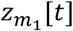 and 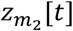 are the analytic representations of rows in ***Y***, i.e., 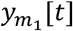 and 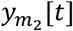, respectively; that is:

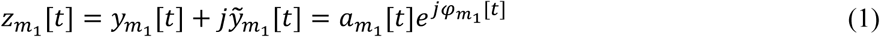

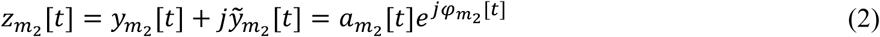

where 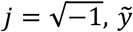 denotes the Hilbert transform of *y*, 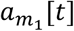 and 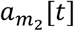 denote instantaneous amplitudes, and 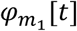 and 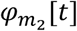 are instantaneous phases of 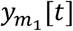 and 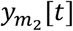. The band-pass filtered fMRI time-series are assumed to be narrow-band satisfying Bedrosian’s theorem (Bedrosian, 1963). The two signals 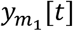 and 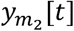 are phase-locked of order 1:1 when:

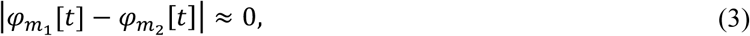

where ∣.∣ is the absolute value operator. We quantified instantaneous phase difference between the signal pairs 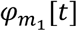 and 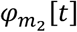 - obtained from all pairs of rows in the analytic associate matrix ***Z*** - as follows:

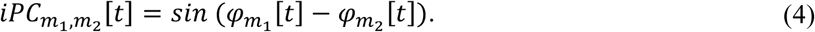

The sinusoid accounts for phase wrapping and ambiguity in the sign of phases over time. Calculation of Eq. (4) for all pairs of nodes and all-time points yields a three-dimensional matrix 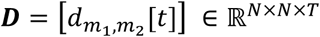 where *T = 1200* for the HPC data and *T = 200* for the validation data. At each time point, we computed the phase coherence quantity (1 − *iPC*) between signal pairs, ranging from 0 (no phase coherence) to 1 (maximal phase coherence).

### 2.5 Correlation-based sliding-windows analysis (CSWA)

CSWA was estimated by calculating the Pearson’s correlation coefficient (*r*) between all pairs of nodes with a single TR window overlap. The Pearson’s correlation coefficient between two time-series,*X*[*t*]and *Y*[*t*] can be written as:

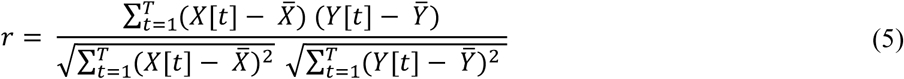

where 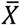 and 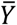 denote sample means and *r* ranges from −1 (full anti-correlation) to 1 (full correlation).

In this study, window lengths varied from 2 TR’s (1.44 seconds) to 250 TR’s (180 seconds) for the HPC data and 2 TR’s (6 seconds) to 60 TR’s (180 seconds) for the validation data. For HCP data, window length increments were set to 2 TR’s resulting in 125 separate window lengths. For validation data, window length increments were set to a single TR resulting in 60 separate window lengths.

### 2.6 Hamming windowing

In our primary analysis, we tapered correlation-based windows (i.e., CSWA) using a Hamming function. We noted that the correlation between IPSA and CSWA weakened when the Hamming function was not used in CSWA (see Supplementary materials 2). This effect is likely to be less of a concern for longer correlation windows than used in our primary analysis (i.e., in ∼60-120 second windows instead of ∼20 second windows) as they will be temporally smoother.

### 2.7 Alignment of IPSA and CSWA time-series

We temporally aligned IPSA and CSWA time-series by assigning the central time point of each CSWA window to the time point of the corresponding IPSA signal. We calculated the cross-correlation between IPSA and CSWA time-series to make sure that the maximal association between the two signals occurred at the central time-point of CSWA. When we overlapped IPSA and CSWA time-series with a rectangular window of *W* time points, *W*/2 time points at the beginning and end of the IPSA signal were disregarded because CSWA time-series always have fewer time-points than IPSA time-series (see the ‘flat’ edges of the CSWA signal in Fig. 4A and B). This ensured that the potential of spectral leakage in IPSA was minimized.

### 2.8 Simulating temporally related time-series using Cholesky decomposition

We also simulated 38503 correlated time-series each associated with 1200 time-points using the Cholesky decomposition algorithm. This is the same number of time-series and time-points as our empirical analysis: 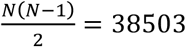 and *T* = 1200. The purpose of this simulation was to further understand the inherent mathematical relationship between IPSA and CSWA, and to assess whether their association was similar between well controlled simulation data (Cholesky decomposition algorithm) and our empirical results (fMRI connectivity).

Let ***C*** ∈ ℝ^n×n^ be a desired correlation matrix and ***X*** ∈ ℝ^n×T^ be a matrix consisting of *n* random and uncorrelated time-series of length *T*. Here, we assume that the matrix ***C*** is real, Hermitian and positive definite. Using Cholesky decomposition, we can factorize ***C*** into the form of ***C*** = ***LL***^′^where ***L*** is a lower-triangle matrix and ′ denotes the transpose operator. The product of ***Y*** = ***LX*** gives a new set of *n* random time-series of covariance matrix ***C*** (i.e., a set of pre-defined correlation values). Across all connection-pairs, ***C*** have correlation values between −1 and 1.

### 2.9 Quantifying the relationship between IPSA and CSWA

#### 2.9.1 Similarity between IPSA and CSWA time-series

We quantified the similarity between IPSA and CSWA time-series using the Spearman’s correlation coefficient measure. This measure accounts for potential monotonic (but not necessarily linear) associations between IPSA and CSWA. This was performed for both original CSWA values (referred to as *preserving negative correlations*) and absolute CSWA values (referred to as *removing negative correlations).*

#### 2.9.2 The effects frequency intervals and negative correlations have on the IPSA/CSWA relationship

We used Cohen’s *d* and 95^th^ percentile confidence intervals to quantify changes in the relationship between IPSA and CSWA as a result of band-pass filtering and negative correlations. Note that negative correlations refer to functional connectivity and not the similarity between IPSA and CSWA.

#### 2.9.3 Time-averaged correlations between ISPA and CSWA brain nodes

We quantified the topological association between IPSA and CSWA nodes by calculating Spearman’s rho and non-linear fitting (6^th^ polynomial order) between time-averaged IPSA and CSWA node matrices that were averaged across all subjects.

## 3. Results

### 3.1 Whole-brain averaged IPSA and CSWA are temporally related

We observed a clear temporal relationship between whole-brain averaged IPSA and CSWA time-series when calculating absolute values of CSWA (Fig. 2 A and D). The strongest correlation between IPSA and CSWA occurred at a window length of 19 second (TR = 24) for the HCP data and 21 seconds (TR = 7) for the validation data (Fig. 2 A and D - black vertical lines). Narrow-band frequency fMRI data (0.03-0.07Hz – Fig. 2 blue colour) exhibited stronger correlations between IPSA and CSWA time-series than wide-band frequency fMRI data (0.01-0.1Hz – Fig. 2 red colour). As seen in Fig. 2 C and F, subjects with many negative correlations in CSWA time-series showed a proportionally weaker association between IPSA and CSWA (while preserving negative correlations). See Table 1 for detailed summary statistics of this data. The remainder of our analyses were conducted at the CSWA length strongest correlated with IPSA (19 seconds for HCP data and 21 seconds for validation data).

**Figure 2:**
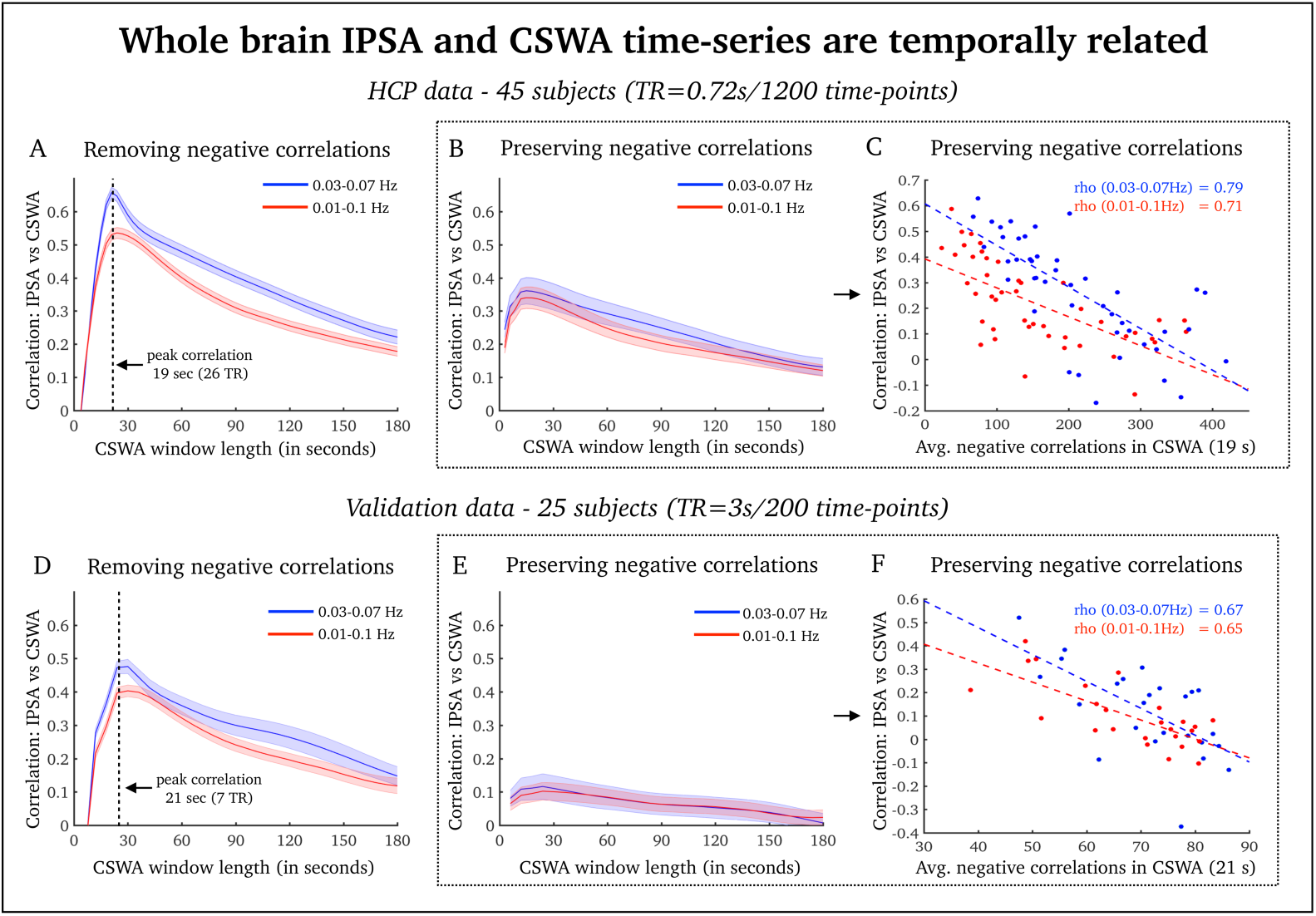
Here we provide an overview of the association between whole brain averaged IPSA and CSWA. **A and D)** HCP data (A) and validation data (D) removing negative correlation in CSWA. **B and E)** HCP data (B) and validation data (E) preserving negative correlation in CSWA. **C and F)** The association between average number of time-points with correlation values below 0 and corresponding Spearman’s rho correlation values between IPSA and CSWA. Number of negative correlations were averaged across all 278 nodes. Each dot represents a single subject. Blue = relationship between IPSA and CSWA for 0.03-0.07Hz; red = relationship between IPSA and CSWA for 0.01-0.1Hz. Shaded areas = standard error of the group. Black stippled lines = sliding window length associated with the strongest correlation between whole brain averaged IPSA and CSWA time-series.

**Table 1:** Summary statistics: IPSA vs CSWA whole-brain time-series averaged across all window lengths. Cohen’s d: ∼0.2: small effect size, ∼0.5: medium effect size, ∼0.8: large effect size.

| **IPSA/CSWA: effect of frequency bands** | **Cohen's $d$** | **CI 95%** | **df** | **p-value** |
| --- | --- | --- | --- | --- |
| HCP data: 0.03-0.07Hz vs. 0.01-0.1Hz<br>(preserving negative correlations) | 0.16 | -0.26-0.58 | 88 | 0.442 |
| HCP data: 0.03-0.07Hz vs. 0.01-0.1Hz<br>(removing negative correlations) | 0.69 | 0.25-1.11 | 88 | 0.002 |
| Validation data: 0.03-0.07Hz vs. 0.01-0.1Hz<br>(preserving negative correlations) | 0.05 | -0.51-0.61 | 48 | 0.849 |
| Validation data: 0.03-0.07Hz vs. 0.01-0.1Hz<br>(removing negative correlations) | 0.50 | 0.08-1.06 | 48 | 0.079 |
| **IPSA/CSWA: effect of negative correlations** | **Cohen's $d$** | **CI 95%** | **df** | **p-value** |
| HCP data: removing vs preserving negative correlations<br>(0.03-0.07Hz) | 0.88 | 0.43-1.30 | 88 | <0.001 |
| HCP data: removing vs preserving negative correlations<br>(0.01-0.1Hz) | 0.82 | 0.38-1.25 | 88 | <0.001 |
| Validation data: removing vs preserving negative correlations<br>(0.03-0.07Hz) | 1.66 | 0.98-2.29 | 48 | <0.001 |
| Validation data: removing vs preserving negative correlations<br>(0.01-0.1Hz) | 1.85 | 1.15-2.50 | 48 | <0.001 |

### 3.2 IPSA and CSWA are temporally related in pair-wise comparison of nodes

There was also a strong temporal association between IPSA and CSWA connection pairs when we removed negative correlations in CSWA, at a correlation window length of ∼20 seconds (Fig. 3 – bars on the right). This relationship was consistent across subjects with low intra-subject variance (Spearman’s rho values ∼0.8-0.9 for HCP data - Fig. 3A right). There were also significantly higher correlation values between IPSA and CSWA in the narrow band-pass filtered fMRI data compared to wider band-pass filtered fMRI data. See Table 1 for detailed summary statistics.

**Figure 3:**
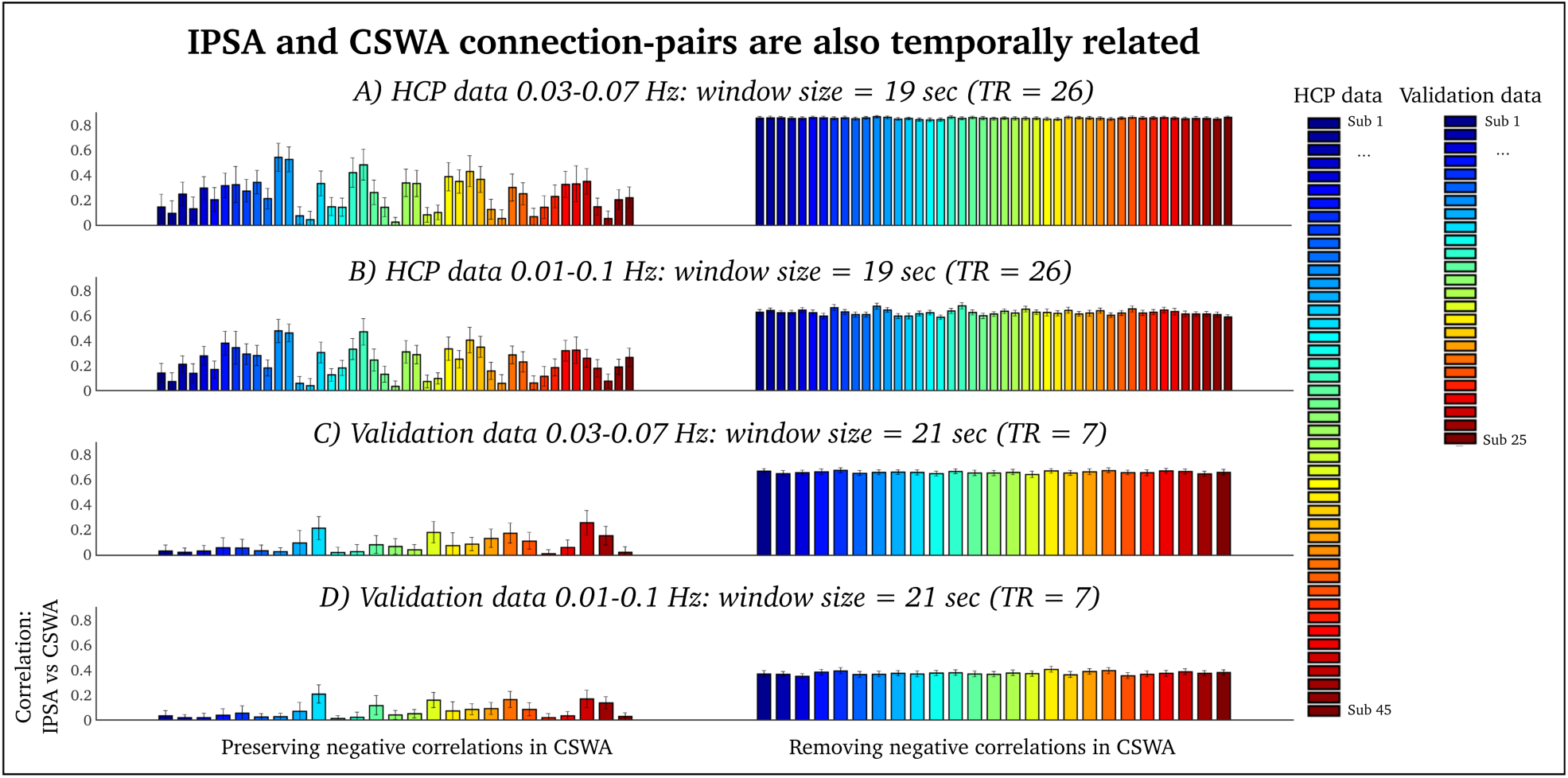
IPSA and CSWA node-pairs are also temporally related. Here we provide Spearman’s rho correlations between all possible IPSA and CSWA connection pairs for each subject (N(N-1)/2=38503 connection pairs). **A)** HCP data - 0.03-0.07Hz. **B)** HCP data - 0.01-0.1Hz. **C)** Validation data - 0.03-0.07Hz. **D)** Validation data - 0.01-0.1Hz. On the left side of the figure we present subject-specific results while preserving negative correlations in CSWA. On the right side of the figure we present subject-specific results after removing negative correlations in CSWA. Error bars = standard deviation.

In Fig. 4, we display IPSA and CSWA time-series of two different connection pairs. These two time-series highlight the effect of calculating absolute CSWA values. In particular, the increase in similarity between IPSA and CSWA connection pairs after calculating absolute correlation is evident in the upper time-series of Fig. 4 A.

**Figure 4:**
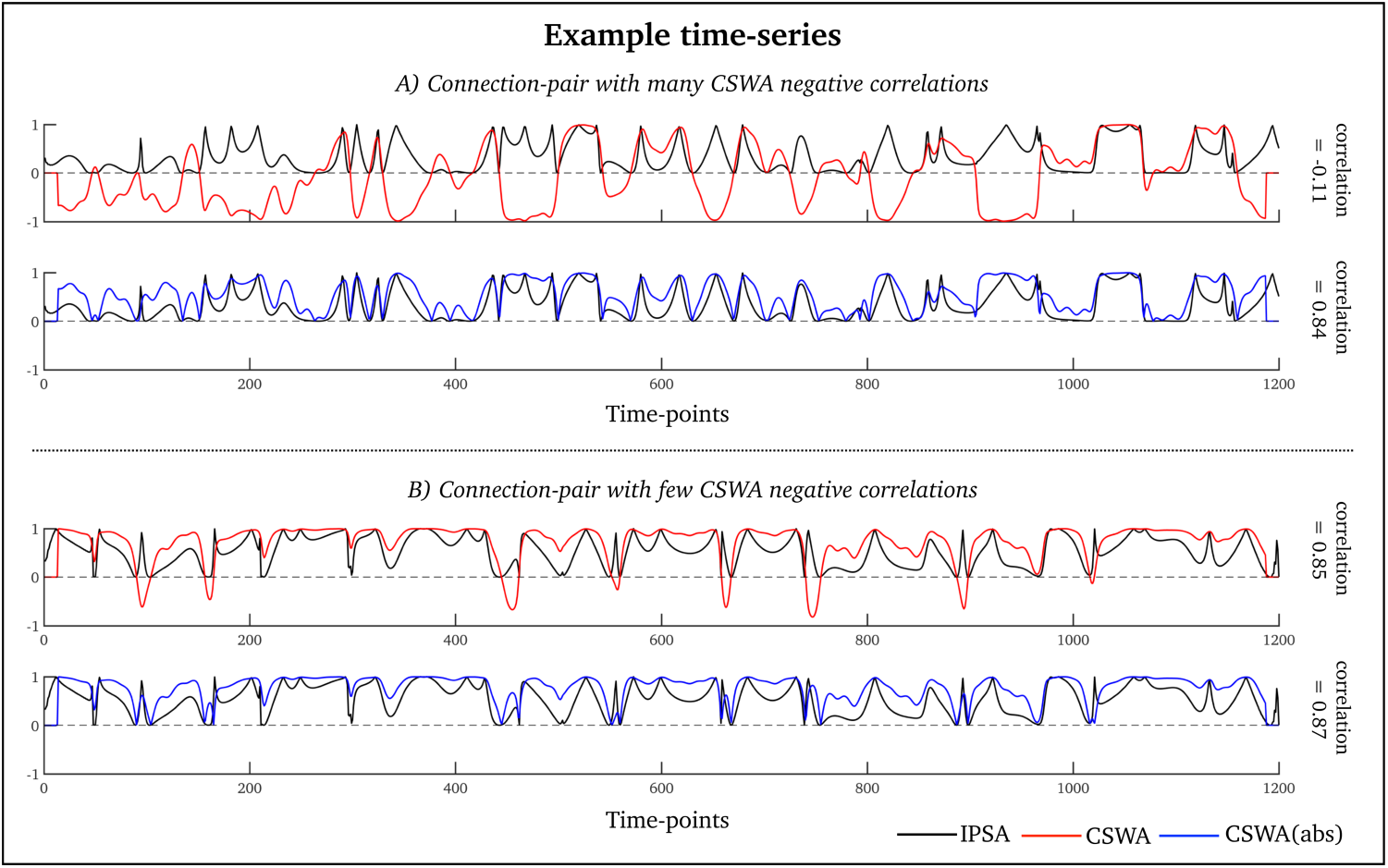
Here we display time-series associated with two connection pairs from the HCP data at 0.03-0.07Hz. **A)** A connection-pair with many negative correlations in CSWA. **B)** A connection pair with few negative correlations in CSWA. Black = IPSA. Red = CSWA. Blue = absolute CSWA.

### 3.3 IPSA and CSWA nodes are topologically related

Lastly, we estimated the topological relationship between specific IPSA and CSWA – i.e., the correlation between time-averaged and group averaged IPSA and CSWA node matrices. Time-averaged IPSA and CSWA matrices were non-linearly related in both the HCP and validation group (scatter plots in Fig. 5 A-D display average IPSA and CSWA values across all subjects), and was similar to the association between pairwise fMRI phase locking values and Pearson’s correlation coefficients as reported by Ponce-Alvarez et al. (2015). The topological relationship between IPSA and CSWA was stronger for HCP data (group averaged Spearman’s rho values = ∼0.8-0.9) compared to the validation data (group averaged Spearman’s rho values = ∼0.4-0.5).

**Figure 5:**
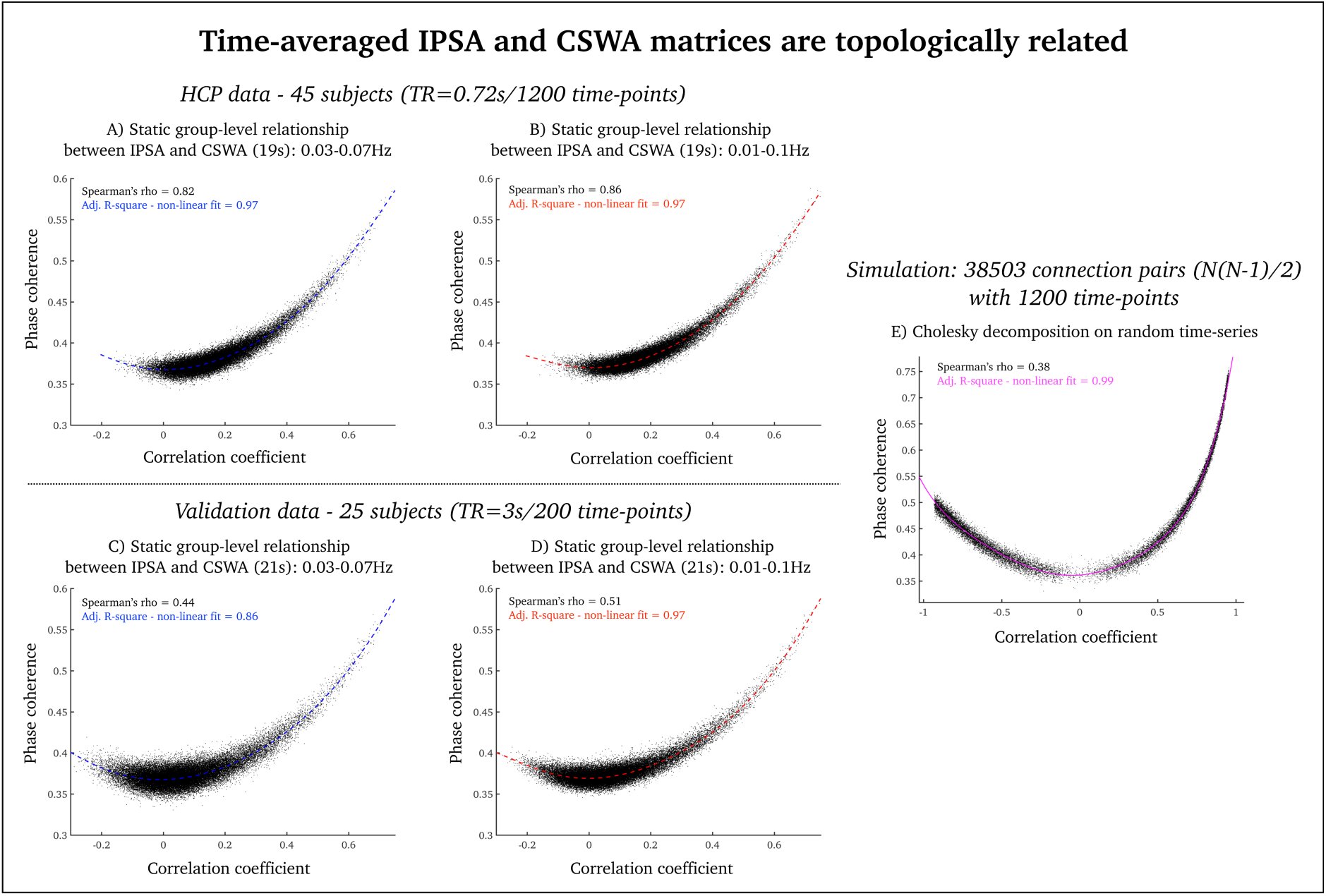
Topological association between IPSA and CSWA: **A and B)** scatter plot displaying the time-averaged correlation between IPSA and CSWA brain nodes, averaged over all subjects for HCP data and 0.03-0.07Hz (A) and 0.01-0.1Hz (B). **C and D)** scatter plot displaying the time-averaged correlation between IPSA and CSWA brain nodes, averaged over all subjects for validation data and 0.03-0.07Hz (C) and 0.01-0.1Hz (D). **E)** Here, we simulated the relationship between IPSA and CSWA using the Cholesky factorisation procedure. Stippled coloured lines denote the best non-linear fit line using the 6th polynomial order of the data.

By simulating correlated time-series based on a Cholesky decomposition, we showed that the non-linear relationship between phase coherence and correlation coefficient is not specific to fMRI data but it rather reflects inherent mathematical features of IPSA and CSWA (Fig 5 E).

The non-linear topological correlation between IPSA and CSWA can theoretically be linearized by calculating absolute values to CSWA in addition to a Fisher’s *z* variance-stabilizing transformation of the CSWA and IPSA data (Fisher, 1915 – see supplementary materials 3).

**Table 2:** Summary statistics: IPSA vs CSWA - connection pairs at peak window length. Cohen’s d: ∼0.2: small effect size, ∼0.5: medium effect size, ∼0.8: large effect size.

| **IPSA/CSWA: effect of frequency bands** | **Cohen's $d$** | **CI 95%** | **df** | **p-value** |
| --- | --- | --- | --- | --- |
| HCP data: 0.03-0.07Hz vs. 0.01-0.1Hz<br>(preserving negative correlations) | 0.16 | -0.25-0.57 | 88 | 0.446 |
| HCP data: 0.03-0.07Hz vs. 0.01-0.1Hz<br>(removing negative correlations) | 15.5 | 13.0-17.7 | 88 | <0.001 |
| Validation data: 0.03-0.07Hz vs. 0.01-0.1Hz<br>(preserving negative correlations) | 0.22 | -0.34-0.78 | 48 | 0.431 |
| Validation data: 0.03-0.07Hz vs. 0.01-0.1Hz<br>(removing negative correlations) | 26.5 | 20.7-31.4 | 48 | <0.001 |
| **IPSA/CSWA: effect of negative correlations** | **Cohen's $d$** | **CI 95%** | **df** | **p-value** |
| HCP data: removing vs preserving negative correlations<br>(0.03-0.07Hz) | 6.38 | 5.29-7.36 | 88 | <0.001 |
| HCP data: removing vs preserving negative correlations<br>(0.01-0.1Hz) | 4.57 | 3.73-5.32 | 88 | <0.001 |
| Validation data: removing vs preserving negative correlations<br>(0.03-0.07Hz) | 11.8 | 9.21-14.1 | 48 | <0.001 |
| Validation data: removing vs preserving negative correlations<br>(0.01-0.1Hz) | 7.28 | 5.59-8.73 | 48 | <0.001 |

## 4. Discussion

In this paper, we systematically quantified the temporal and topological association between two different measures of time-resolved fMRI connectivity, namely IPSA and CSWA. We provide converging evidence that IPSA and CSWA most likely convey much of the same information as we observed a strong temporal and topological relationship between IPSA and absolute CSWA. This was consistent in 70 subjects across two different datasets and is in line with a previous study by Glerean et al. (2012) where it was also reported that fMRI based IPSA and CSWA time-series are temporally related. Our results both replicates and builds on this work in several ways. Firstly, we highlight the importance of window length in correlation based fMRI analysis. Secondly, we demonstrate the effect that negative correlations (in CSWA) have on the relationship between IPSA and CSWA. Thirdly, we outline the significance of narrow-band pass filtering in phase synchrony analyses of fMRI data. Lastly, we quantify the time-averaged and topological association between IPSA and CSWA. Next, we provide an in-depth discussion of these points.

### 4.1 IPSA versus CSWA: correlation window length, repetition time and frequency interval

One of the main goal of this study was to quantify whether fMRI window length, band-pass filtering and/or sampling rate affect the relationship between IPSA and CSWA. We found that the correlation between IPSA and CSWA was highly dependent on the length of sliding windows. Both of our fMRI datasets had a peak correlation between IPSA and CSWA at window lengths of approximately twenty seconds, unrelatedly to the underlying fMRI sampling rate. Although the temporal similarity between IPSA and CSWA time-series appears to be independent from fMRI sampling rate, we do not imply that higher fMRI sampling rate is not beneficial for time-resolved fMRI connectivity analysis. Statistically speaking we kept our two dataset as separate entities, but there were stronger correlations between IPSA and CSWA time-series in the fMRI data with high sampling rate (Fig. 2 and 3). Also, the relationship between time-averaged IPSA and CSWA matrices were stronger for HCP data compared to validation data (Fig. 5). This is not surprising given that ∼1200 time-points were averaged for HCP data compared to only ∼200 time-points for validation data. Thus, faster sampling rate (which results in a high number of sample points) is likely to improve the statistical evaluation of IPSA and CSWA, but it is unlikely to affect the slow frequency fluctuations underlying time-resolved fMRI connectivity.

We also observed that the correlation between IPSA and CSWA was stronger for narrow-band frequency data compared to wider-band frequency data. This is a finding that is consistent with the Bedrosian’s requirements for phase synchrony data (Bedrosian, 1963) and highlight the significance of narrow-band pass filtering for this type of analysis (Omidvarnia et al 2016). It still remains to be seen whether 0.03-0.07Hz is the optimal narrow frequency interval for IPSA. It was outside the scope of the current study to test more than two frequency band between IPSA and CSWA because of the already exhaustive computational and interpretational complexities of our analyses.

### 4.2 A note on ‘anti-correlations’

The relationship between IPSA and CSWA is highly dependent on negative correlations in CSWA because strong ‘anti-correlations’ (in CSWA) have high phase coherence (in IPSA) (see Fig. 4A). After we enforced absolute values upon CSWA, we observed that the correlation between IPSA and CSWA increased drastically (Fig. 2 and 3). While this reinforces that these two methods most likely convey comparable types of information it will have interpretational consequences for both temporal and topological analyses. For example, in graph theoretic analysis of fMRI it is common to threshold connectivity matrices by retaining only the strongest connected nodes in the graph. For phase synchrony analysis, such a thresholding procedure will inevitably include ‘anti-corrrelated’ node pairs in the graph. Although negative correlations are likely to be biologically informative (Chen et al., 2011; Zhan et al., 2017), we recommend that care needs to be taken when comparing and interpreting graph theoretical measures derived from thresholded CSWA and IPSA matrices.

### 4.3 Future direction and limitation

It is a contemporary and contentious question to ask whether time-resolved fMRI connectivity display non-stationarity behaviour (see Hindriks et al., 2016; Laumann et al., 2016.; Zalesky et al., 2014). In other words, *is time-resolved fMRI connectivity more informative than time-averaged fMRI connectivity?* Although the purpose of this paper was not to directly deal with this issue, we want to raise a potentially noteworthy point regarding instantaneous phase synchrony and non-stationarity. Non-stationarity in a CSWA time-series can be (theoretically) detected when strong positive correlations temporally transit towards strong negative correlations, or vice versa. Such temporal transitions may go undetected in phase synchrony signals since strong negative and positive correlations will both have high phase synchrony (see Fig. 4). In our future work, we aim to assess how IPSA perform in comparison to CSWA using non-parametric null models of stationarity, including Vector Autoregressive Null Models (Sims, 1980) and Phase Randomisation procedures that preserves the correlational structure of the data (Prichard and Theiler, 1994).

A limitation of this study (and time-resolved fMRI connectivity studies in general) was our inability to quantify whether IPSA is an inherently ‘better’ or ‘worse’ measure of time-resolved fMRI connectivity than CSWA. This is because it is still relatively uncertain what constitutes ‘good’ and ‘bad’ time-resolved fMRI connectivity in terms of signal variance. We expect to grasp more of an understanding about the fundamental quality of these measures as time-resolved fMRI connectivity research continues to evolve.

## 5. Conclusion

Our findings suggest that IPSA and CSWA convey comparable information of time-resolved fMRI connectivity. Although IPSA require narrow-band fMRI filtering, we are inclined to favour IPSA over CSWA as there is no need for selecting appropriate window length and overlap. A MATLAB code for calculating IPSA is provided in Supplementary materials 4.

## Acknowledgements

The primary fMRI data in this study was provided by the Human Connectome Project, WUMinn Consortium (1U54MH091657; Principal Investigators: David Van Essen and Kamil Ugurbil) funded by the 16 National Institutes of Health (NIH) institutes and centers that support the NIH Blueprint for Neuroscience Research; and by the McDonnell Center for Systems Neuroscience at Washington University. We thank Mira Semmelroch and Magdalena Kowalczyk for acquisition of the fMRI validation data. This study was supported by the National Health and Medical Research Council (NHMRC) of Australia (#628952). The Florey Institute of Neuroscience and Mental Health acknowledges the strong support from the Victorian Government and in particular the funding from the Operational Infrastructure Support Grant. We also acknowledge the facilities, and the scientific and technical assistance of the National Imaging Facility (NIF) at the Florey node and The Victorian Biomedical Imaging Capability (VBIC). GJ is supported by an NHMRC practitioner’s fellowship (#1060312).

## References

Allen, E.A., Damaraju, E., Plis, S.M., Erhardt, E.B., Eichele, T., Calhoun, V.D., 2012. Tracking Whole-Brain Connectivity Dynamics in the Resting State. Cereb. Cortex 24, 663–676.

Ashburner, J., 2007. A fast diffeomorphic image registration algorithm. NeuroImage 38, 95–113.

Bedrosian, E., 1963. A product theorem for Hilbert transforms. Proc. IEEE 51, 868–869.

Behzadi, Y., Restom, K., Liau, J., Liu, T.T., 2007. A component based noise correction method (CompCor) for BOLD and perfusion based fMRI. NeuroImage 37, 90–101.

Biswal, B., Yetkin, F.Z., Haughton, V.M., Hyde, J.S., 1995. Functional connectivity in the motor cortex of resting human brain using echo-planar MRI. Magn. Reson. Med. 34, 537–541.

Bullmore, E., Sporns, O., 2009. Complex brain networks: graph theoretical analysis of structural and functional systems. Nat. Rev. Neurosci. 10, 186–198.

Chang, C., Glover, G.H., 2010. Time–frequency dynamics of resting-state brain connectivity measured with fMRI. NeuroImage 50, 81–98.

Chen, G., Chen, G., Xie, C., Li, S.-J., 2011. Negative Functional Connectivity and Its Dependence on the Shortest Path Length of Positive Network in the Resting-State Human Brain. Brain Connect. 1, 195– 206.

Córdova-Palomera, A., Tornador, C., Falcón, C., Bargalló, N., Brambilla, P., Crespo-Facorro, B., Deco, G., Fañanás, L., 2016. Environmental factors linked to depression vulnerability are associated with altered cerebellar resting-state synchronization. Sci. Rep. 6, 37384.

Cribben, I., Wager, T.D., Lindquist, M.A., 2013. Detecting functional connectivity change points for single-subject fMRI data. Front. Comput. Neurosci. 7, 143.

Demirtaş, M., Tornador, C., Falcón, C., López-Solà, M., Hernández-Ribas, R., Pujol, J., Menchón, J.M., Ritter, P., Cardoner, N., Soriano-Mas, C., Deco, G., 2016. Dynamic functional connectivity reveals altered variability in functional connectivity among patients with major depressive disorder. Hum. Brain Mapp. 37, 2918–2930.

Fisher, R.A., 1915. Frequency Distribution of the Values of the Correlation Coefficient in Samples from an Indefinitely Large Population. Biometrika 10, 507–521.

Fornito, A., Zalesky, A., Breakspear, M., 2015. The connectomics of brain disorders. Nat. Rev. Neurosci. 16, 159–172.

Fransson, P., 2005. Spontaneous low-frequency BOLD signal fluctuations: an fMRI investigation of the resting-state default mode of brain function hypothesis. Hum. Brain Mapp. 26, 15–29.

Friston, K.J., Ashburner, J.T., Kiebel, S.J., Nichols, T.E., Penny, W.D., 2011. Statistical Parametric Mapping: The Analysis of Functional Brain Images: The Analysis of Functional Brain Images. Academic Press.

Friston, K.J., Williams, S., Howard, R., Frackowiak, R.S., Turner, R., 1996. Movement-related effects in fMRI time-series. Magn. Reson. Med. 35, 346–355.

Glasser, M.F., Sotiropoulos, S.N., Wilson, J.A., Coalson, T.S., Fischl, B., Andersson, J.L., Xu, J., Jbabdi, S., Webster, M., Polimeni, J.R., Van Essen, D.C., Jenkinson, M., 2013. The minimal preprocessing pipelines for the Human Connectome Project. NeuroImage 80, 105–124.

Glerean, E., Salmi, J., Lahnakoski, J.M., Jääskeläinen, I.P., Sams, M., 2012. Functional Magnetic Resonance Imaging Phase Synchronization as a Measure of Dynamic Functional Connectivity. Brain Connect. 2, 91–101.

Hindriks, R., Adhikari, M.H., Murayama, Y., Ganzetti, M., Mantini, D., Logothetis, N.K., Deco, G., 2016. Can sliding-window correlations reveal dynamic functional connectivity in resting-state fMRI? NeuroImage 127, 242–256.

Hudetz, A.G., Liu, X., Pillay, S., 2015. Dynamic repertoire of intrinsic brain states is reduced in propofol-induced unconsciousness. Brain Connect. 5, 10–22.

Hutchison, R.M., Womelsdorf, T., Allen, E.A., Bandettini, P.A., Calhoun, V.D., Corbetta, M., Della Penna, S., Duyn, J.H., Glover, G.H., Gonzalez-Castillo, J., Handwerker, D.A., Keilholz, S., Kiviniemi, V., Leopold, D.A., de Pasquale, F., Sporns, O., Walter, M., Chang, C., 2013. Dynamic functional connectivity: promise, issues, and interpretations. NeuroImage 80, 360–378.

Kaiser, R.H., Whitfield-Gabrieli, S., Dillon, D.G., Goer, F., Beltzer, M., Minkel, J., Smoski, M., Dichter, G., Pizzagalli, D.A., 2016. Dynamic Resting-State Functional Connectivity in Major Depression. Neuropsychopharmacology 41, 1822–1830.

Laumann, T.O., Snyder, A.Z., Mitra, A., Gordon, E.M., Gratton, C., Adeyemo, B., Gilmore, A.W., Nelson, S.M., Berg, J.J., Greene, D.J., McCarthy, J.E., Tagliazucchi, E., Laufs, H., Schlaggar, B.L., Dosenbach, N.U.F., Petersen, S.E., n.d. On the Stability of BOLD fMRI Correlations. Cereb. Cortex 1–14.

Lindquist, M.A., Xu, Y., Nebel, M.B., Caffo, B.S., 2014. Evaluating dynamic bivariate correlations in resting-state fMRI: a comparison study and a new approach. NeuroImage 101, 531–546.

Mormann, F., Lehnertz, K., David, P., E. Elger, C., 2000. Mean phase coherence as a measure for phase synchronization and its application to the EEG of epilepsy patients. Phys. Nonlinear Phenom. 144, 358–369.

Omidvarnia, A., Pedersen, M., Walz, J.M., Vaughan, D.N., Abbott, D.F., Jackson, G.D., 2016. Dynamic regional phase synchrony (DRePS): An Instantaneous Measure of Local fMRI Connectivity Within Spatially Clustered Brain Areas. Hum. Brain Mapp. 37, 1970–1985.

Pedersen, M., Omidvarnia, A., Curwood, E.K., Walz, J.M., Rayner, G., Jackson, G.D., 2017a. The dynamics of functional connectivity in neocortical focal epilepsy. NeuroImage Clin. 15, 209–214.

Pedersen, M., Omidvarnia, A., Walz, J.M., Zalesky, A., Jackson, G.D., 2017b. Spontaneous brain network activity: Analysis of its temporal complexity. Netw. Neurosci. 1, 100–115.

Ponce-Alvarez, A., Deco, G., Hagmann, P., Romani, G.L., Mantini, D., Corbetta, M., 2015. Resting-State Temporal Synchronization Networks Emerge from Connectivity Topology and Heterogeneity. PLoS Comput Biol 11, e1004100.

Preti, M.G., Bolton, T.A., Van De Ville, D., 2016. The dynamic functional connectome: State-of-the-art and perspectives. NeuroImage. In press.

Prichard, D., Theiler, J., 1994. Generating surrogate data for time series with several simultaneously measured variables. Phys. Rev. Lett. 73, 951–954.

Rashid, B., Arbabshirani, M.R., Damaraju, E., Cetin, M.S., Miller, R., Pearlson, G.D., Calhoun, V.D., 2016. Classification of schizophrenia and bipolar patients using static and dynamic resting-state fMRI brain connectivity. NeuroImage 134, 645–657.

Shakil, S., Lee, C.-H., Keilholz, S.D., 2016. Evaluation of sliding window correlation performance for characterizing dynamic functional connectivity and brain states. NeuroImage 133, 111–128.

Shen, X., Tokoglu, F., Papademetris, X., Constable, R.T., 2013. Groupwise whole-brain parcellation from resting-state fMRI data for network node identification. NeuroImage 82, 403–415.

Sims, C.A., 1980. Macroeconomics and Reality. Econometrica 48, 1–48.

Smith, S.M., 2002. Fast robust automated brain extraction. Hum. Brain Mapp. 17, 143–155.

Smith, S.M., Miller, K.L., Moeller, S., Xu, J., Auerbach, E.J., Woolrich, M.W., Beckmann, C.F., Jenkinson, M., Andersson, J., Glasser, M.F., Essen, D.C.V., Feinberg, D.A., Yacoub, E.S., Ugurbil, K., 2012. Temporally-independent functional modes of spontaneous brain activity. Proc. Natl. Acad. Sci. 109, 3131–3136.

Tagliazucchi, E., Laufs, H., 2014. Decoding Wakefulness Levels from Typical fMRI Resting-State Data Reveals Reliable Drifts between Wakefulness and Sleep. Neuron 82, 695–708.

Thompson, W.H., Fransson, P., 2015. The frequency dimension of fMRI dynamic connectivity: Network connectivity, functional hubs and integration in the resting brain. NeuroImage 121, 227–242.

Tzourio-Mazoyer, N., Landeau, B., Papathanassiou, D., Crivello, F., Etard, O., Delcroix, N., Mazoyer, B., Joliot, M., 2002. Automated anatomical labeling of activations in SPM using a macroscopic anatomical parcellation of the MNI MRI single-subject brain. NeuroImage 15, 273–289.

Van Essen, D.C., Smith, S.M., Barch, D.M., Behrens, T.E.J., Yacoub, E., Ugurbil, K., 2013. The WU-Minn Human Connectome Project: An overview. NeuroImage 80, 62–79.

Yu, Q., Erhardt, E.B., Sui, J., Du, Y., He, H., Hjelm, D., Cetin, M.S., Rachakonda, S., Miller, R.L., Pearlson, G., Calhoun, V.D., 2015. Assessing dynamic brain graphs of time-varying connectivity in fMRI data: Application to healthy controls and patients with schizophrenia. NeuroImage 107, 345–355.

Zalesky, A., Fornito, A., Cocchi, L., Gollo, L.L., Breakspear, M., 2014. Time-resolved resting-state brain networks. Proc. Natl. Acad. Sci. 111, 10341–10346.

Zhan, L., Jenkins, L.M., Wolfson, O., GadElkarim, J.J., Nocito, K., Thompson, P.M., Ajilore, O., Chung, M.K., Leow, A., 2017. The Significance of Negative Correlations in Brain Connectivity. J. Comp. Neurol. 525, 3251–3265.

